# Genome-wide characterization of extant clonal diversity in Chilean Carménère

**DOI:** 10.64898/2026.04.03.716224

**Authors:** Jadran Garcia, Noé Cochetel, Jimena Balic, Samuel Barros, Rosa Figueroa-Balderas, Alvaro Castro, Dario Cantu

**Affiliations:** Department of Viticulture and Enology, University of California Davis, Davis, CA 95616, USA; UC Davis-Chile Life Sciences Innovation Center, Santiago 7520424, Chile; UNIVIVEROS, Paine, Chile; Genome Center, University of California Davis, Davis, CA 95616, USA

**Keywords:** Carménère, Clonal variation, Whole-genome resequencing, Replicate-aware variant calling, Grape genomics, Somatic mutations, Genetic resources

## Abstract

Carménère is a widely cultivated and internationally recognized grapevine cultivar in Chile, yet genetic variation among its clones remains poorly characterized. Early studies based on SSR and AFLP markers detected limited polymorphism, but these approaches interrogate only a small fraction of the genome, leaving the extent of clonal diversity unresolved. Here, we generated an improved chromosome-scale diploid genome assembly of Carménère FPS clone 02 and characterized clonal genomic diversity by sequencing 36 biological replicates representing 12 clones maintained in Chile, including heritage selections rescued from old producer vineyards by Viña Santa Carolina as part of its Bloque Herencia conservation program, and commercial nursery-derived clones. Focusing on low-frequency variants and using replicate-aware consensus calling, we identified more than 9,000 private single nucleotide variants (SNVs) and small indels per clone, providing high-resolution markers for clonal identification. Although most variants were located in repetitive or intergenic regions, a subset affected coding sequences, with genes involved in plant-pathogen interactions, transport, and secondary metabolism most frequently impacted. While variant-affected genes associated with wine anthocyanin content, TA, pH, and alcohol percentage were identified, broader phenotypic characterization will be required to assess their biological significance. Overall, this study provides a genome-wide characterization of extant clonal diversity in Carménère, with implications for clonal selection and genetic resource conservation.

## Introduction

Carménère is a red-skinned wine grape cultivar (*Vitis vinifera* L.) that was historically cultivated in France but largely eliminated following the phylloxera epidemic of the late 19th century (Banerjee et al., 2010; Belancic & Agosin, 2007; kenwytsma, 2023; Pszczólkowski T., 2004). Its susceptibility to coulure, late-ripening phenology, and relatively low yields compared to cultivars such as Merlot made it a poor candidate for replanting, leading to its near disappearance from European vineyards by the early 20th century, despite its later preservation and expansion elsewhere (Belancic & Agosin, 2007; Pszczólkowski T., 2004; Santos-Buch, 2019).

In the 1990s, a grape widely cultivated in Chile and marketed as Merlot was identified as Carménère. This misidentification was first noted through field observations by French ampelographer Jean-Michel Boursiquot in 1994 and later confirmed using molecular markers (Hinrichsen et al., 2001), demonstrating that Carménère had been present in Chile for more than a century (Pszczólkowski T., 2004). Chile’s geographic isolation, which prevented the introduction of phylloxera, allowed ungrafted, pre-epidemic plant material to persist, and Carménère proved well adapted to local climates and soils (Ceballos, 2014; Müller, 2004). Officially recognized as a distinct cultivar in 1998, it has since emerged as Chile’s flagship red variety, with more than 10,000 hectares planted nationwide (OIV, 2017; Palma, 2024; SAG, 2021).

Carménère in Chile originates from cuttings introduced from France in the mid-19th century, capturing only a limited fraction of the genetic diversity present in France at the time (Pszczólkowski T., 2004). Subsequent propagation relied largely on this founding material, with limited additional input from French and Italian nurseries. This narrow genetic base, combined with more than a century of vegetative propagation without systematic clonal selection, raises critical questions about the extent and structure of clonal diversity available to Chilean viticulture.

Early efforts to characterize Carménère diversity relied on SSR and AFLP markers applied to 26 accessions from Chile, France, and Italy, and reported extremely limited polymorphism: only two of 20 SSR loci were variable, and just seven of the 344 fragments detected by the AFLP analysis were polymorphic (Moncada & Hinrichsen, 2007). However, these marker systems interrogate a small fraction of the genome and are largely insensitive to the point mutations and small indels that accumulate during vegetative propagation, likely leading to a substantial underestimation of clonal diversity (Fischer et al., 2017). By contrast, whole-genome resequencing approaches provide much higher resolution. In a study of 46 grapevine clones across four cultivars, mapping reads to cultivar-specific reference genomes revealed on average ∼4 million variants per clone, and subsequent filtering identified markers capable of discriminating individual clones within each cultivar (Urra et al., 2023).

Applying these approaches to Carménère requires high-quality genomic resources. The first Carménère genome assembly, generated using long read sequencing, captured structural and nucleotide level variation between homologous chromosomes but lacked chromosome scale scaffolding, limiting its utility for fine scale analyses of genome structure and variant phasing (Minio et al., 2019). To overcome this limitation, we generated a chromosome scale diploid reference genome and sequenced the whole genomes of 12 Carménère clones in triplicate, for a total of 36 samples, to resolve clonal genetic diversity in Chile. By focusing on low frequency, replicate-supported variants, we identified more than 9,000 clone private single nucleotide variants (SNVs) and small insertion and deletions (INDELs) per clone, providing high resolution markers for clonal identification. We further show that these variants preferentially affect genes associated with plant-pathogen interaction, transport, and secondary metabolism.

## Materials and methods

### Plant material origin and preparation

The Carménère clones used in this study include two selections. The first group consists of seven clonal selections developed by Viña Santa Carolina (VSC) through its ‘Bloque Herencia’ project, launched in 2011 to rescue and characterize pre-phylloxera grapevine material from old Chilean vineyards. Each selection originated from a single outstanding mother plant identified in long-established vineyards across historically significant viticultural zones: Cauquenes (Maule Valley), Puente Alto (Maipo Valley), Almahue (Cachapoal Valley), and Melozal (Loncomilla Valley). Mother plants were rescued between 2011 and 2015 and established at the Bloque Herencia experimental block at Fundo Totihue (Requinoa, Cachapoal Valley) between 2013 and 2016, where they have since been maintained and characterized agronomically, phenotypically, and enologically. The second group comprises nursery-derived clones from commercial programs developed jointly by AgroUC-Copec and Pontificia Universidad Católica de Chile (PUC), selected from traditional Chilean vineyards planted with material imported from France around 1870 and evaluated over eight seasons in a clonal selection block in San Fernando. Two additional French selections imported to Chile in 2009 through a commercial nursery in the Maipo Valley were also included. Clone origins and additional details are provided in **Supplementary Table 1**.

Tissue samples for DNA extraction were collected from actively growing shoots (young leaves) of three biological replicates per selection. High-molecular-weight genomic DNA (gDNA) was extracted from young leaves using the method described by (Chin et al., 2016). DNA purity was assessed with a Nanodrop 2000 spectrophotometer (Thermo Scientific, IL, USA), and DNA concentration was quantified using the DNA High Sensitivity Kit on a Qubit 2.0 Fluorometer (Life Technologies, CA, USA).

DNA sequencing libraries were prepared using 1 μg of the extracted genomic DNA, following the Kapa LTP Library Preparation Kit protocol (Kapa Biosystems, MA, USA). Library quantity and quality were evaluated using the High Sensitivity chip on a Bioanalyzer 2100 (Agilent Technologies, CA, USA). Sequencing was performed on an Illumina HiSeq platform (Novogene Co. Ltd., Beijing, China) using a 150 bp paired-end (PE) read configuration.

HiFi sequencing libraries were prepared from the material previously collected in Minio (2019) following the manufacturer’s protocol for the SMRTbell Express Template Prep Kit 3.0 (Pacific Biosciences, CA, USA). Libraries were size-selected using the LightBench (Yourgene Health, FL, USA) and subsequently sequenced on a PacBio Revio platform at the DNA Technology Core Facility, University of California, Davis, CA, USA.

### Carménère FPS 02 genome assembly and annotation

The genome assembly was generated from HiFi reads (Yield: 27.62 Gb) by combining the Hifiasm v0.19.8-r603 (Cheng et al., 2021) outputs with a high-density gene map (Cochetel et al., 2025) using HaploSync v1.0 (Minio et al., 2022). After testing multiple configurations, the parameters selected for hifiasm assembly that ensure the best balance between haplotype for size, marker and BUSCO model distribution were: “-f 25 -s 0.25 -D 3 -n 11”. Markers were located in each haplotype using minimap2 (Li, 2021) and the parameters: “-x splice:hq --cs=long -G 30000 -u b”. BUSCO v6.0.0 (Tegenfeldt et al., 2025) was used with the lineage dataset eudicotyledons_odb12. RepeatMasker v4.1.7-p1 (Smit et al., 2015) was used for the annotation of the repeats.

Telomeric repeat sequences were identified independently for each Carménère haplotype using TIDK v0.2.63 (Telomere Identification Toolkit; (Brown et al., 2025). The tidk search module was run with the sequence TTTAGGG and a window size of 10 kb.

Gene models were annotated using a weighted, multiple evidence strategy as proposed in (Heuermann et al., 2025) and applied in (Baptiste, 2025) with 3 independent annotation tools. First, the reference genome and gene models of PN40024 v5.1 were obtained from Grapedia (https://grapedia.org/). These gene models were transferred onto the Carménère FPS 02 assembly (haplotype 1, haplotype 2, and unplaced contigs) using Liftoff v1.6.3 (Shumate & Salzberg, 2021), with the options “-copies, - polish, and -cds”.

Next, Braker was run on Carménère FPS 02 using RNAseq data and a database of proteins. The RNAseq data was obtained from the bioproject PRJNA1244791 (Cochetel et al., 2025) which included leaves and berries of Cabernet Franc, Cabernet Sauvignon and Sauvignon Blanc at different physiological stages. The protein database was the Viridiplantae clade of the OrthoDB v.11 (https://bioinf.uni-greifswald.de/bioinf/partitioned_odb11/) and the set of proteins from PN40024 v5.1. BRAKER3 (Gabriel et al., 2024) executed using a Singularity container built from the Docker image teambraker/braker3:latest (build date: 7 November 2025; Ubuntu 22.04 base), run with Apptainer version 1.3.2 (Kurtzer et al., 2021).

Then, gene models of Carménère FPS 02 also independently predicted using Helixer v0.3.6 (Holst et al., 2025) executed within a Singularity container built from the Docker image gglyptodon/helixer-docker:helixer_v0.3.6_cuda_12.2.2-cudnn8 (build date: 12 November 2025; Ubuntu 22.04 base), and run using Apptainer version 1.3.2.

The resulting annotation of each of the 3 methods were cleaned by removing gene models for which all isoforms were classified as non-coding, as well as removing individual non-coding isoforms from genes that also contained at least one valid protein-coding isoform. Then the tool agat_sp_complement_annotations.pl v1.4.1 (Jacques Dainat et al., 2026) was used to combine the annotation using as base the lifted PN40024 v5.1 models, giving it the highest weight, followed by the braker models and last the Helixer models. One extra round of non-coding isoforms cleaning was made in order to secure pseudogenes in the final set.

A BUSCO v6.0.0 (Tegenfeldt et al., 2025) analysis with the database Eudicotyledons OrthoDB v12 dataset was made to evaluate the annotation completeness.

### Analysis synteny and conservation of gene models of PN40024 v5.1 in the new assembly

Synteny relationships between the Carménère genome and the PN40024 v5.1 reference genome were identified using a protein-based collinearity approach. The longest isoform per gene per genome were retained and combined (Haplotype 1 + PN40024 v5.1 and Haplotype 2 + PN40024 v5.1). All-vs-all sequence similarity searches were performed using DIAMOND blastP v2.1.9.163 (Buchfink et al., 2021) with an e-value threshold of 1e-10, in sensitive mode, retaining up to 100 target sequences per query. Next, gene coordinates were extracted from GFF3 of each combination and concatenated. Collinearity blocks were then identified using MCScanX v1.0 (Wang et al., 2012) with default parameters.

To quantify cross-genome synteny, collinearity outputs were parsed to extract gene pairs between PN40024 and Carménère haplotypes. Redundant matches were removed, and unique syntenic gene pairs were retained for downstream analyses. The proportion of genes in syntenic blocks was calculated for both genomes based on these gene pairs.

To complement the synteny analysis, gene content was compared between PN40024 v5.1 and the Carménère genome was assessed using GMAP. Genome databases were built independently for each Carménère haplotype and unplaced contigs, as well as for the PN40024 v5.1 reference. Coding sequences (CDS) from PN40024 v5.1 were mapped to each Carménère assembly using GMAP v2024-11-20 (Wu & Watanabe, 2005) with parameters allowing up to five alignments per query and multithreaded execution. Alignments were filtered to retain transcripts with ≥80% sequence identity and ≥80% coverage. Genomic coordinates of retained alignments were converted into BED format and intersected with Carménère gene annotations using BEDTools intersect v2.29.1 (Quinlan, 2014), requiring at least 80% overlap. The proportion of PN40024 v5.1 transcripts supported by Carménère gene annotations was calculated for each haplotype and across the combined assembly including the unplaced.

### Structural variation between haplotypes and hemizygosity calculation

Structural variation between Carménère haplotype 1 and haplotype 2 was identified using a whole-genome alignment approach. Soft-masked haplotype assemblies aligned using Minimap2 v 2.22-r1101 (Li, 2021) in assembly-to-assembly mode with options -x asm5 and --eqx. The resulting alignments were sorted and indexed using SAMtools. Structural variants were then detected using SyRI v1.6.3 (Goel et al., 2019). To focus on larger sequence variants, insertions, deletions, were filtered to retain events greater than 50 bp. Variant counts and cumulative lengths were calculated per category to summarize structural divergence between haplotypes.

Additionally, gene-level hemizygosity between haplotypes was assessed by reciprocal mapping of coding sequences (CDS) using GMAP v2024-11-20. CDS sequences from each haplotype annotation were aligned against the opposite haplotype genome using GMAP with parameters allowing multiple alignments and multithreading. Alignments were filtered to retain transcripts with ≥80% identity and ≥80% coverage. The proportion of hemizygous genes was calculated based on the total number of unmapped gene set between the two haplotypes.

### Functional annotation

A deep functional annotation was performed using a combination of sequence similarity searches and domain-based annotation. Protein sequences from the Carménère genome were compared against the NCBI RefSeq plant protein database (ftp://ftp.ncbi.nlm.nih.gov/refseq, retrieved January 17th, 2017) using DIAMOND blastP v2.1.9.163 with an e-value threshold of 1e-3, retaining up to 100 target sequences per query. BLAST results were imported into Blast2GO for functional annotation, where Gene Ontology (GO) terms were assigned based on sequence similarity. To complement similarity-based annotation, protein domain and functional signatures were identified using InterProScan, and associated GO terms were integrated into the annotation dataset. Within Blast2GO, the combined annotation set was enrichment using ANNEX, enzyme code assignment, KEGG pathway mapping, and GO-slim classification using the plant-specific GO subset. The final functional annotation set combined sequence similarity, domain-based evidence, and curated ontology mappings.

To focus on pathway and network associated with grapevine, an homology annotation was also made following the VitisNet v12X (Grimplet et al., 2009, 2012). This was performed by identifying the best PN40024 v5.1 homolog for each Carménère gene. Because VitisNet v12X annotations are available for PN40024-derived gene models, Carménère haplotypes were processed independently to establish correspondence between Carménère and PN40024 transcripts. First, a DIAMOND blastP v2.1.9.163 (Buchfink et al., 2021) in all-versus-all mode was applied to the proteins of PN40024 and each Carménère haplotype with sensitive settings (e-value ≤ 1e-10, maximum 100 target sequences). Second, transcript coordinates were extracted from GFF3 annotations and used with MCScanX v1.0.0 (Wang et al., 2012) to infer collinear relationships between PN40024 and each Carménère haplotype. Syntenic orthologs were prioritized as the primary homolog assignment, and the best collinear match per Carménère transcript was retained. For Carménère genes not assigned through synteny, the best PN40024 protein hit from blastP, excluding self-hits, was used as a secondary homolog assignment. The combined set of synteny-based and similarity-based homologs was used to transfer PN40024 functional labels and VitisNet network annotations to Carménère genes.

### Clone sequencing and variant calling

The sequencing data of the 36 samples were quality and adapter trimmed using fastp v.1.0.1 (Chen, 2025) with the options “-detect_adapter_for_pe -q 20 -l 100”. The trimmed reads were subsampled to get 20X haploid coverage of the newly assembled genome of Carménère FPS 02. The resulting reads were processed using NVIDIA Parabricks (pbrun fq2bam) version 4.5.0-1 (Zhu et al., 2025), run in the Docker container nvcr.io/nvidia/clara/clara-parabricks:4.5.0-1 with GPU acceleration. For each sample, paired-end reads were aligned independently to each Carménère FPS 02 haplotype using the Parabricks implementation of BWA-MEM (Li, 2013), followed by coordinate sorting and duplicate marking. GPU-based sorting and BAM writing were enabled (--gpusort, --gpuwrite). Variant calling was carried out using NVIDIA Parabricks HaplotypeCaller using the same version as described above. For each sample, variants were called independently in GVCF mode against each haplotype of Carménère FPS 02. Variant calling parameters included a minimum base quality score of 20 and a minimum confidence threshold for calling variants of 10.

Per-sample GVCF files generated with Parabricks HaplotypeCaller were combined using GATK CombineGVCFs (GATK v4.5.0.0; McKenna et al., 2010) and subsequently jointly genotyped with GATK GenotypeGVCFs, using each haplotype of Carménère FPS 02. Both steps were executed within the Docker container broadinstitute/gatk:4.5.0.0.

Jointly genotyped variant call sets were post-processed separately for each haplotype. Variants were first left-normalized and decomposed into biallelic records using bcftools norm (v1.8-30-gb717d08; Danecek et al., 2021), with respect to the corresponding haplotype reference genome. SNVs and INDELs were then separated using GATK SelectVariants (GATK v4.5.0.0). Variant-level hard filtering was applied independently to SNPs and INDELs using GATK VariantFiltration. SNPs were filtered based on quality by depth (QD < 2.0), strand bias (FS > 60.0; SOR > 3.0), mapping quality (MQ < 40.0), and rank sum tests (MQRankSum < −12.5; ReadPosRankSum < −8.0). INDELs were filtered using thresholds for QD < 2.0, FS > 200.0, SOR > 10.0, and ReadPosRankSum < −20.0. Variants failing any filter were flagged and excluded from downstream analyses.

### Clonal relatedness analysis

For the clonal exploratory relatedness analysis, variants were additionally filtered to remove sites with excessive missing genotypes. Specifically, sites with missing genotype calls in ≥20% of samples were excluded using bcftools view with the expression COUNT(GT=“mis”)/N_SAMPLES < 0.2. A 20% missingness threshold was chosen as a conservative compromise to retain informative sites while minimizing missing-data artifacts in pairwise relatedness and clustering analyses. This filter was applied to the VCF from the previous step, focusing on SNPs sets for each Carménère haplotype. The filtered SNP datasets were converted to PLINK2 (Chang et al., 2015) binary format using --make-bed. Pairwise kinship coefficients were estimated using the KING-robust implementation in PLINK2 (--make-king-table). Kinship matrices were generated for each haplotype-specific SNP dataset and used for downstream clustering and ordination analyses.

Kinship matrixes were imported into R for visualization and exploratory analyses. Heatmaps were produced using ComplexHeatmap (Gu, 2022), with hierarchical clustering based on Euclidean distances. For ordination, PCA was performed using prcomp() on the kinship matrix, and MDS was computed using classical multidimensional scaling (cmdscale) on a distance transformation (1 − kinship). Based on the results the sample Carolina77_RF20 was removed.

### Clone consensus variants

To focus downstream analyses on true clonal variation, low-frequency variants, consistent with somatic variation, were filtered based on alternative allele frequency. First, only the Carménère samples were retained and variant sites with missing genotypes across the retained samples were removed. Then, alternative allele count (AC), allele number (AN), and alternative allele frequency (AF) annotations were computed for each haplotype and variant type (SNV and INDELs) using bcftools +fill-tags. Variants were then filtered to retain low-frequency alternative alleles (AF ≤ 0.25) using bcftools view. This threshold was chosen to reduce the contribution of widely shared or fixed polymorphisms likely reflecting ancestral or cultivar-level variation, while retaining rare and low-frequency variants informative for clonal differentiation. To assess whether variants shared among samples were enriched for intra-clone sharing, we compared the observed proportion of intra-clone variants to a null expectation based on random sampling of 3 individuals. The expected intra-clone probability was calculated using combinatorial probabilities. Enrichment was quantified as the ratio of observed to expected intra-clone proportions. Statistical significance was evaluated using one-sided Fisher’s exact test.

### Variant effect annotation

To locate the clonal variants within the genomic regions of the Carménère assembly, the previously filtered variants were intersected with different annotated genomic features. Repetitive regions were sorted and merged to produce a non-redundant repeat interval set in BED format. The intervals corresponding exons and introns were extracted from the gene annotation and converted to BED format. Overlapping intervals within each feature class were merged to avoid redundant counting. Intergenic regions were defined as the genomic space between gene intervals across, and the intergenic regions without repeats was generated by subtracting the repetitive elements intervals. Variant coordinates passing allele-frequency and replicate-consensus filters were converted to BED format and intersected with each genomic feature class using BEDTools intersect v2.29.1 (Quinlan, 2014). Intersections were performed separately for each haplotype and variant type (SNPs and INDELs). Both variant-level annotations (retaining clone-specific consensus flags) and summarized count tables were generated to quantify the distribution of variants across genomic features. Feature lengths were calculated from merged BED intervals and used for normalization.

Functional effects of filtered variants were annotated using SnpEff v5.1 (Cingolani, 2022) with the Carménère FPS 02 reference genome. Separate SnpEff databases were constructed for each haplotype using the annotated gene models (GTF file, and corresponding genomic sequences, CDS, and protein sequences) and variant effect annotation was performed independently for SNPs and INDELs. Upstream and downstream regions extending 3 kb from annotated transcripts were included. For each variant, predicted functional effects, affected genes, and impact categories were reported. Annotated variants were subsequently parsed to summarize effect classes and impact categories at both the variant and gene levels. Variants annotated exclusively as “MODIFIER” were excluded from downstream impact-focused analyses, allowing emphasis on low-, moderate-, and high-impact effects.

### Phenotypic correlation

To identify potential candidate genes associated with phenotypic variation among Carménère clones, we performed an exploratory phenotype-consistency analysis using clone-level variant calls and available phenotypic data. This analysis is based on the assumption that rare variants will be enriched among individuals at the tails of the phenotypic distribution (Amanat et al., 2020). Since this study only looked into a small number of samples, variants were aggregate into gene-burden style carrier/non-carrier status per gene to increase detection power (Floriani & Lipka, 2025). With this approach, instead of testing individual variants, multiple variants within a gene are aggregated and the association is studied at the gene level with gene carrying and not carrying variants. Only Chilean clones with available phenotypic information were included (**Supplementary Table 1**). Variants classified as consensus heterozygous within each clone and predicted to have high or moderate functional impact were retained for the analysis.

Continuous phenotypic traits were analyzed by defining extreme groups for each trait, using the three clones with the lowest and the three clones with the highest observed values. For each impacted gene, clone-level carrier matrices were constructed to indicate the presence or absence of at least one qualifying variant (high- or moderate-impact). These matrices were then compared against the trait-defined high and low groups to quantify phenotype consistency. For each impacted gene-trait combination, we calculated the proportion of carrier clones in the high and low groups, the difference in carrier frequency between groups, and a consistency score based on the number of carrier and non-carrier clones matching either the high- or low-trait model. Genes were then ranked according to consistency score, and those showing perfect association (defined as complete agreement between all carrier/non-carrier clones and all the clones belonging to either the high or low trait group) were retained for further evaluation.

Given the limited number of clones and incomplete phenotypic coverage across traits, this analysis should be interpreted cautiously and considered an exploratory framework for identifying potential candidate genes and pathways putatively associated with phenotypic divergence among clones. The R code used to perform the analysis can be found in (https://github.com/jadgarci/Carmenere-diversity-supporting-scripts)

## Results

### A chromosome-scale assembly of Carménère FPS 02

To investigate genetic variation among Carménère clones, a high-quality reference genome of Carménère FPS 02 was first assembled from PacBio HiFi long reads at 55x haploid coverage. Hifiasm contigs (Cheng et al., 2021) were scaffolded and phased with a genetic map using HaploSync (Minio et al., 2022), resulting in a chromosome-scale diploid assembly, with each haplotype comprising 19 chromosomes and a size of 479.6 ± 1.7 Mbp, for a total genome size of 959.3 Mbp placed in phased chromosomes (**Table 1**). Unplaced contigs accounted for only 32.9 Mbp (0.03% of total assembly length). Most telomeric regions were successfully identified across both haplotypes (55 out of 76 expected telomeres; **Supplementary Figure 1**). Genome completeness assessed by BUSCO recovered 99.8% of the universal single-copy orthologs in the eudicotyledons_odb12 database. Compared to the previous Carménère assembly, which consisted of more than 9,000 scaffolds and achieved 93% BUSCO completeness, the new assembly represents a substantial improvement in both contiguity and completeness, with chromosomes assembled end-to-end.

**Table 1.** Statistics of the genome assembly of Carménère FPS 02.

|  | Combined | Haplotype 1 | Haplotype 2 | Unplaced |
| --- | --- | --- | --- | --- |
| Assembly length (Mbp) | 992.1 | 481.3 | 478 | 32.9 |
| Number of sequences | 338 | 19 | 19 | 300 |
| Average length (Mbp) | 2.9 | 25.3 | 25.2 | 0.1 |
| GC content (%) | 35.0 | 34.6 | 34.7 | 44.2 |
| N50 (Mbp) | 24.9 | 25.5 | 24.9 | 0.2 |
| L50 | 18 | 9 | 9 | 42 |
| Number of gene loci | 72,653 | 35,096 | 35,673 | 1,884 |
| Gene content (Mbp) | 341.4 | 167.1 | 171.1 | 3.2 |
| Repeats (Mbp) | 561.5 | 268.2 | 266.1 | 27.2 |
| Repeats (%) | 56.6 | 55.73 | 55.67 | 82.72 |
| Comp. BUSCOs (%) | 99.8 | 99.8 | 99.8 | 1.1 |
| Comp. and s.c. BUSCOs (%) | 0.1 | 98.4 | 97.9 | 1 |
| Comp. and dup. BUSCOs (%) | 99.7 | 1.4 | 1.9 | 0.1 |
| Fragmented BUSCOs | 0.1 | 0.1 | 0.1 | 0.7 |
| Missing BUSCOs (%) | 0.1 | 0.1 | 0.1 | 98.2 |

Repeat and gene annotation identified 267.2 ± 1.1 Mbp of repetitive sequences per haplotype (∼56%) and 169.1 ± 2.0 Mbp of gene space, encompassing 35,384 ± 289 annotated genes per haplotype and an additional 1,884 genes in the unplaced contigs, resulting in 72,653 annotated gene models overall, compared to 70,109 in the previous assembly. Gene distribution across haplotypes is also more balanced in the new genome. While the previous version showed an uneven distribution (56% of genes in the primary assembly and 44% in haplotigs), the new assembly displays a more even proportion, with 48.3% and 49.1% of genes assigned to haplotype 1 and haplotype 2, respectively, and 2.6% located in unplaced contigs. Functional annotation revealed that, on average, 92.9 ± 0.2% of genes per haplotype were annotated with at least one conserved domain (**Supplementary Table 2**), 69.4 ± 0.6% were assigned Gene Ontology terms (**Supplementary Table 2**), and 60.7 ± 0.7% were linked to VitisNet 12X homologs (**Supplementary Table 3**).

Comparative analyses with the PN40024 v5.1 and this new genome showed that most Carménère genes (57.9 ± 0.3%) are retained in collinear genomic context with PN40024 v5.1 genes (50.1 ± 0.7%) when compared with MCScanX. Coding sequence mapping comparisons revealed an even higher level of gene conservation. Using transcript alignment, 96.5 ± 0.1% of PN40024 v5.1 transcripts aligned to each Carménère haplotype with ≥80% identity and coverage, indicating widespread preservation of gene sequences even when local gene order is disrupted. Furthermore, when aligned transcripts were intersected with gene models across both haplotypes and unplaced contigs, approximately 90% of PN40024 v5.1 transcripts were supported by Carménère gene annotations with at least 80% overlap, demonstrating high correspondence between the two annotation sets despite more limited syntenic conservation.

Structural variation between haplotype 1 and haplotype 2 was characterized using SyRI, revealing 27 inversions spanning 13 Mbp and 219 translocations totaling 3.2 Mbp. In addition, 50 INDELs larger than 50 bp were detected, accounting for 131.9 kbp of sequence divergence. These values represent a substantial reduction relative to the previous assembly, which reported more than 6,000 INDELs totaling 15.3 Mbp. Furthermore, reciprocal CDS mapping between haplotypes indicated that 1.98% of genes are hemizygous between haplotypes.

### Whole-genome sequencing and variant calling of Carménère clones

Genomic analyses were conducted on 36 plants representing 12 distinct Carménère clones maintained in Chile. These clones include seven heritage selections rescued from old producer vineyards across Chile and developed by Viña Santa Carolina under its Bloque Herencia conservation program (Balic, unpublished), as well as nursery-derived selections (AgroUC-Copec and Facultad de Agronomía e Ingenería Forestal de la Pontificia Universidad Católica de Chile), collectively representing the breadth of Carménère clonal material currently available in Chile. Each clone was represented by three biological replicates to support clone-level analyses of genetic variation.

Phenotypic observations related to yield (single berry weight and single bunch weight) and wine chemistry from micro fermentations (alcohol content, anthocyanin concentration, titratable acidity and pH) were obtained from some of these clones (**Supplementary Table 1**). These traits showed distinct levels of variability. Alcohol content in wine showed minimal variation (mean = 13.11%, SD = 0.29, CV = 2.2%). In contrast, anthocyanin concentration and single bunch weight displayed substantially higher variability (CV = 20.9% and 36.6%, respectively), suggesting pronounced phenotypic divergence among clones. Intermediate levels of variation were observed for titratable acidity and single berry weight and pH (CV from 7.7% to 14%).

All clones were sequenced using Illumina, yielding an estimated haploid genome coverage of 23.7 ± 0.3x per plant after quality control filtering and adapter trimming. To place Carménère clonal variation in a broader genomic context and support preliminary clonal identity analyses, sequencing data from 13 previously published grapevine cultivars were included as outgroups in selected downstream analyses. These cultivars span a wide range of genetic distances relative to Carménère, from closely related Bordeaux varieties sharing recent common ancestry (Cabernet Franc, Cabernet Sauvignon, Sauvignon Blanc, and Merlot) to more distantly related cultivars (Chardonnay, Riesling, and Zinfandel; **Supplementary Table 4**). To minimize biases associated with sequencing depth, all datasets were subsampled to an estimated haploid genome coverage of 20x prior to analysis.

The subsampled sequencing reads were mapped independently to each haplotype of the Carménère FPS 02 reference genome, followed by joint genotyping across all samples. After variant-level filtering, 14.86 ± 0.02 million SNVs and 3.50 ± 0.001 million small INDELs were identified per haplotype, corresponding to average densities of 31.9 ± 0.6 SNVs/kbp and 7.6 ± 0.02 INDELs/kbp, respectively (**Supplementary table 5**). Variant density was broadly distributed across chromosomes, but also showing regions with higher-than-expected numbers of variants interspersed with segments displaying pronounced depletion. These patterns differed between SNVs and INDELs and varied slightly between haplotypes (**Supplementary Figure 2**).

### Pairwise kinship and sample quality control

To evaluate genetic relatedness among samples and assess clonal identity within the Chilean Carménère collection, we estimated pairwise kinship coefficients from the genome-wide SNP dataset. Most Carménère samples, including Carménère FPS 02 used to generate the reference genome, formed a tight, well-defined cluster clearly separated from other *V. vinifera* cultivars included as outgroups (**Figure 2**). Within this cluster, kinship coefficients among Carménère samples were uniformly high (∼0.46), consistent with clonal identity, whereas values relative to outgroup cultivars declined with genetic distance, reaching near zero or negative values for more distantly related cultivars such as Riesling and Zinfandel, as expected for unrelated individuals. For reference, kinship coefficients of ∼0.25 are expected between first-degree relatives and ∼0.125 between second-degree relatives. Accordingly, Cabernet Franc and Cabernet Sauvignon, which share a known parent-offspring relationship, showed kinship coefficients of ∼0.22, while Zinfandel showed negative values (∼ −0.18) relative to other cultivars. The uniformly high kinship observed among Carménère clones thus exceeds values expected even for full siblings, confirming their clonal relationship. One sample, Carolina77_RF20, exhibited lower kinship (∼0.31) and clustered closer to outgroup cultivars, suggesting possible misidentification or admixture with a non-Carménère genotype. This result was independently supported by principal component analysis (**Supplementary Figure 3**), and Carolina77_RF20 was excluded from downstream analyses, leaving 35 samples representing 12 clones.

### Variant filtering and clonal consensus calling

Following confirmation of clonal identity and exclusion of Carolina77_RF20, the dataset was restricted to Chilean Carménère clones and sites with missing genotypes were removed. An alternative allele-frequency filter (AF ≤ 0.25) was applied to exclude widely shared heterozygous polymorphisms likely reflecting standing cultivar-level variation, thereby enriching for low-frequency variants consistent with recent somatic mutation (**Supplementary Figure 4**). This filtering strategy is supported by haplotype comparisons, as an average of 88.5 ± 0.6% of the excluded variants overlap haplotype-specific differences identified by SyRI. This filtering reduced the dataset from 3,680,561 ± 2,810 SNVs and 616,169 ± 599 INDELs per sample to 123,828 ± 1,355 SNVs and 39,859 ± 265 INDELs per sample (**Supplementary Table 6**).

To characterize clonal diversity, variants were summarized using a replicate-aware consensus framework. For each clone, variants were classified into three confidence tiers based on concordance across the three biological replicates: consensus variants (detected in all three replicates), mixed variants (detected in two of three), and singletons (detected in only one; **Figure 1**). This approach prioritizes high-confidence clonal variants while retaining information on partially supported and low-confidence calls for downstream analyses.

**Figure 1.**
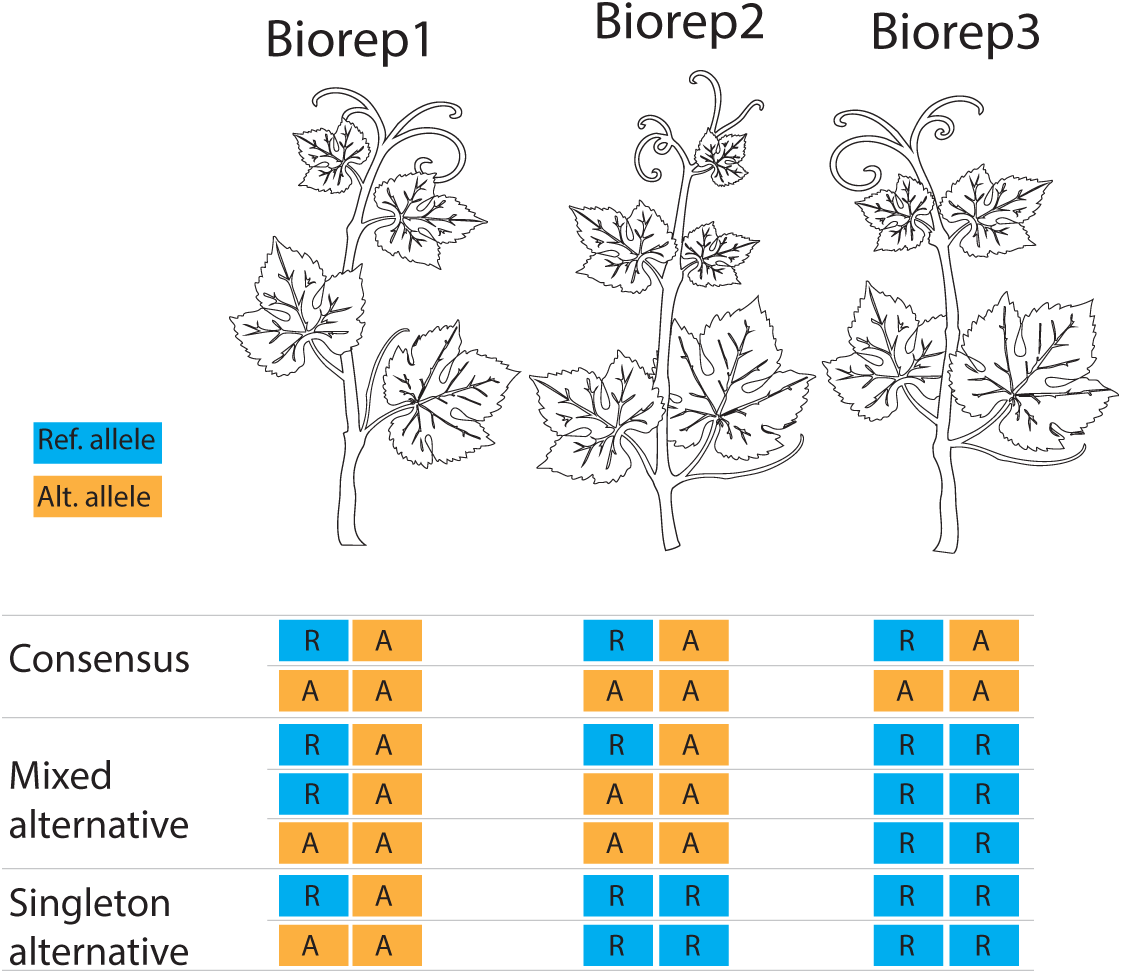
Diagram of the replicate-aware consensus framework used to flagged variants per clone. To derive clone-level consensus genotypes from replicate-based variant calls, we implemented a custom Python workflow that assigns consensus flags to each clone at every variant position based on the genotypes observed across its biological replicates. For each clone, replicate genotypes were evaluated jointly and classified according to predefined consensus (Figure 1). Clones with fully concordant replicate genotypes were assigned unambiguous consensus states (e.g., Consensus heterozygous, Consensus homozygous alternative or Consensus homozygous reference), whereas partial concordance were flagged with graded categories (mixed alternative and single alternative). This approach preserves information on replicate agreement while enabling systematic identification of high-confidence clonal variants calls. The full script and decision logic are available in a publicly accessible repository (https://github.com/jadgarci/Carmenere-diversity-supporting-scripts).

**Figure 2.**
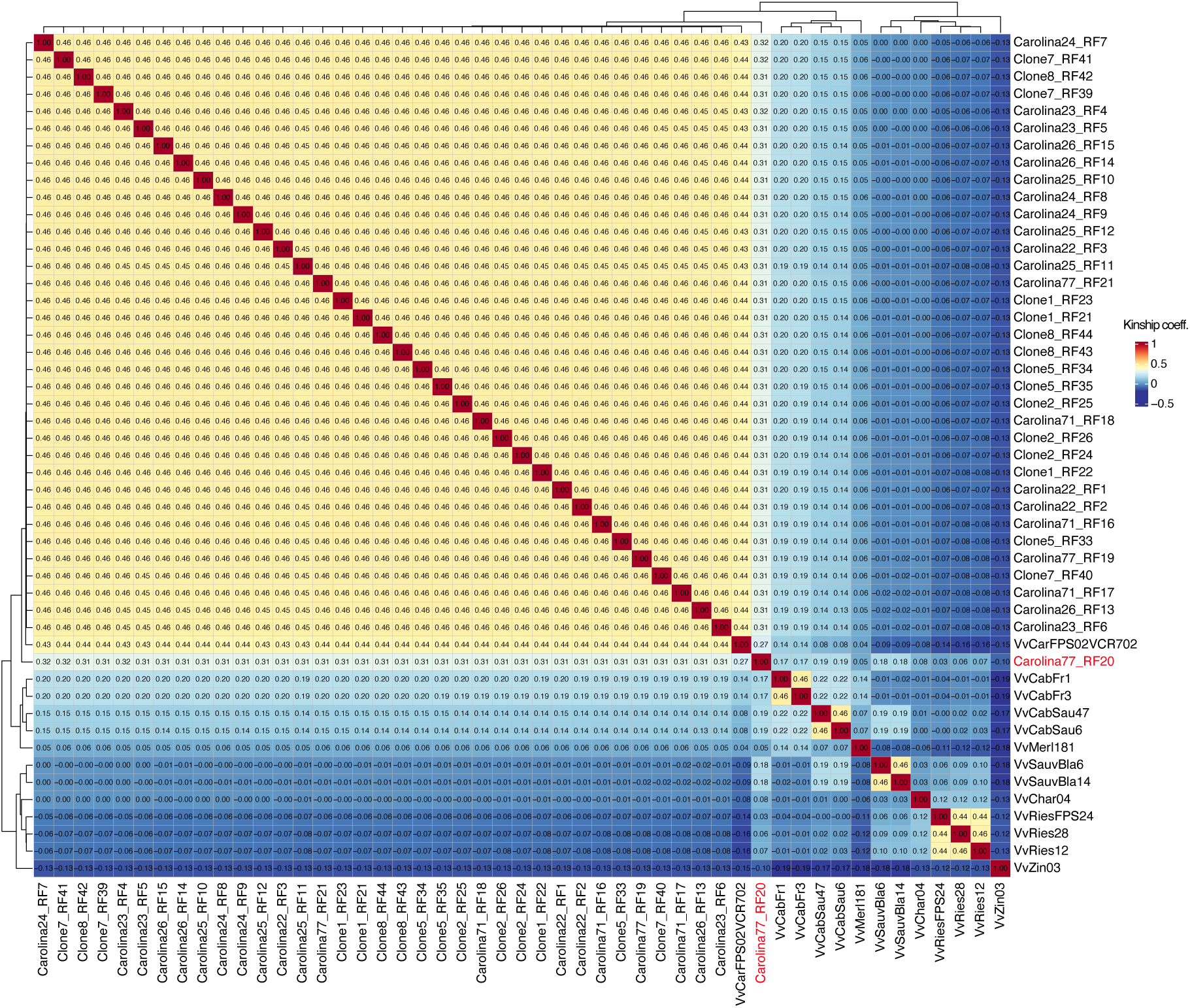
Pairwise kinship heatmap of Carménère clones and outgroup *V. vinifera* cultivars based on genome-wide SNP data from haplotype 1. Kinship coefficients were estimated using PLINK2 and visualized as a clustered heatmap, with color intensity reflecting the magnitude of pairwise relatedness. The sample highlighted in red (Carolina77_RF20) shows consistently lower kinship relative to the other Carménère clones, suggesting possible misidentification or a distinct genetic origin.

The impact of allele-frequency filtering is shown in **Figure 3**. Before filtering, consensus heterozygous variants were largely shared across multiple clones, with counts peaking at 11-12 clones for both SNVs and INDELs across haplotypes. After filtering, the distribution shifted strongly toward clone-private variants (variants present in all replicates of each clone), so when variants were shared by three samples, the likelihood of being three replicates of a clone was higher than any other random samples (**Supplementary Table 7**), suggesting an effective removal of standing cultivar-level variation while retaining low-frequency replicate-supported somatic variants. Most retained consensus heterozygous variants were private to individual clones, with an average of 7,522 ± 1,165 SNVs and 2,045 ± 360 INDELs per clone (**Supplementary Figure 5**; **Additional file 1**). A smaller fraction was shared among up to six clones, consistent with either older somatic mutations predating clonal divergence or shared clonal ancestry within a subset of the collection (**Figure 3**).

**Figure 3.**
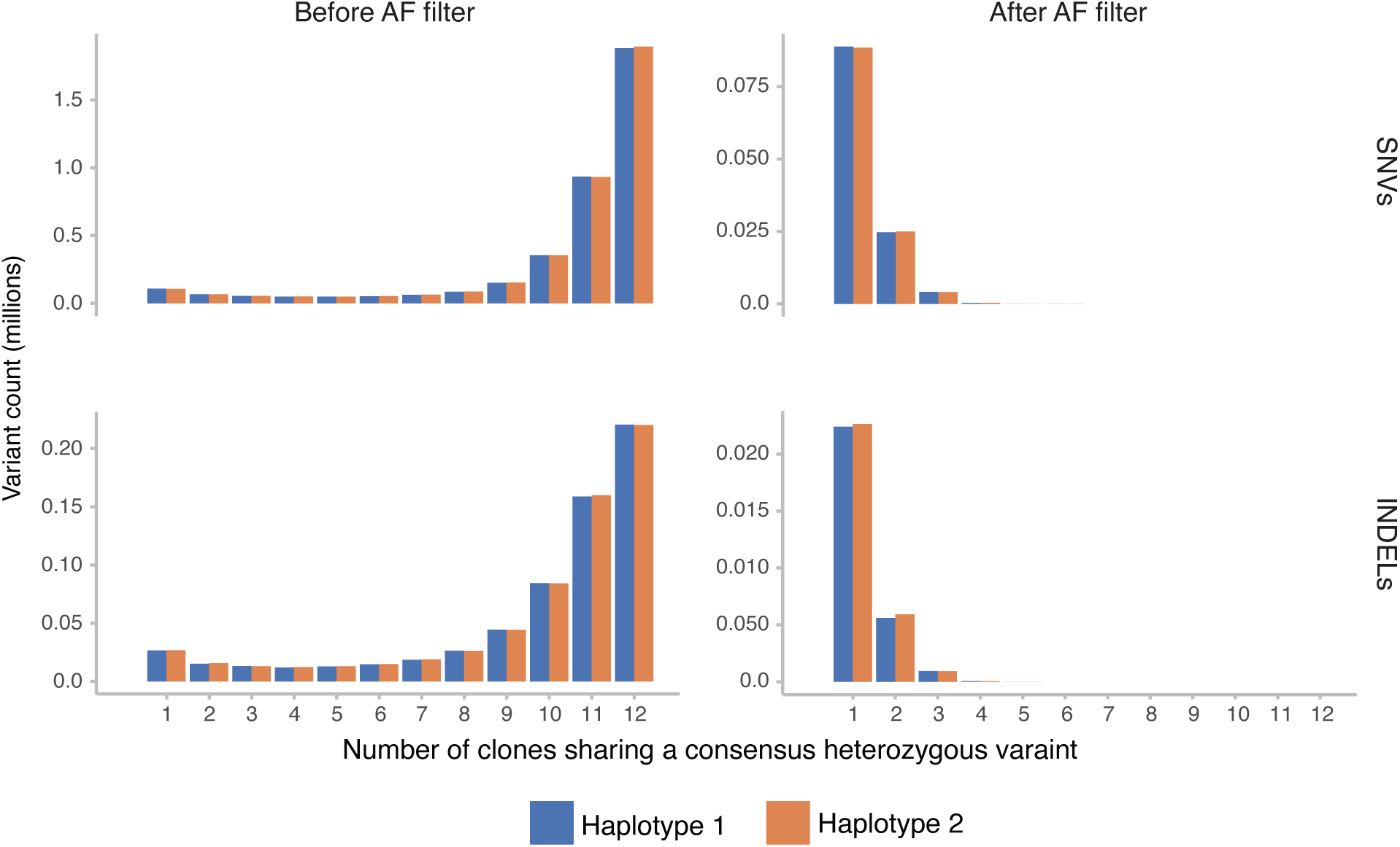
Number of consensus heterozygous variants shared by different numbers of clones. The figure shows the distribution of consensus heterozygous variants SNVs and INDELs observed in exactly k clones, where k ranges from 1 to 12 clones. Distributions are shown before and after applying the alternative allele frequency filter (alternative allele frequency < 0.25).

### Genomic distribution and predicted functional impact of clonal variants

The biological relevance of clonal variation depends on its genomic context and can be inferred by its predicted functional impact. Across both haplotypes and all confidence tiers, variant density was highest within repetitive regions for both SNVs and INDELs (**Figure 4**). Outside repetitive elements, SNV density was broadly similar across exonic, intronic, and intergenic regions, whereas INDEL density varied by feature: intergenic regions showed higher density than introns, and exons exhibited the lowest density, consistent with purifying selection against frameshift mutations in coding sequences. Singleton variants were the most abundant confidence tier across all genomic features, indicating that most detected somatic mutations are private to individual plants and consistent with ongoing mutation accumulation. Variant counts declined steeply with increasing numbers of clones sharing a variant, with very few variants shared among more than three or four clones. Variants located in repetitive regions extended to higher clone-sharing numbers, possibly reflecting elevated mutation rates or older variation predating clonal divergence (**Figure 4**).

**Figure 4.**
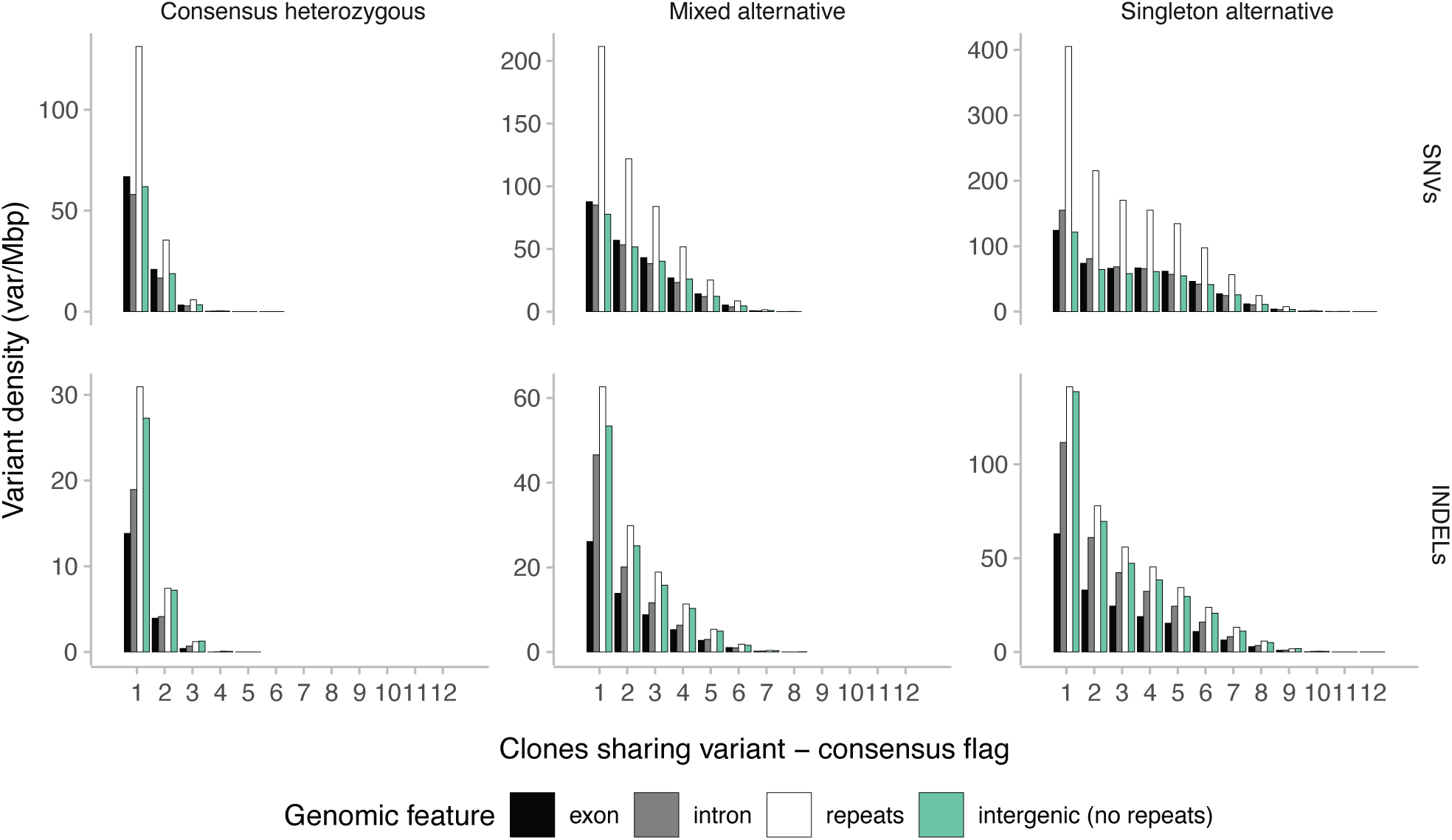
Genomic distribution and density of clonal variants as a function of the number of clones sharing each variant. The figure shows the density of consensus heterozygous, mixed alternative, and singleton variants according to the number of clones carrying each variant. Densities are shown for exonic and intronic regions, annotated repeats, and intergenic regions excluding repeats.

The predicted functional impact of variants was assessed using SnpEff (**Figure 5**). Per haplotype, 8,838 ± 186 variants were classified as high impact, 28,484 ± 459 as moderate impact, and 14,749 ± 234 as low impact, collectively affecting 7,292 ± 169 genes (**Additional file 2; Supplementary Table 8**). Among affected genes, the largest fraction carried only moderate-impact variants (∼30-36%), followed by genes carrying only low-impact variants (∼15-18%), while genes affected exclusively by high-impact variants represented a small minority (∼2%). Approximately 13% of affected genes harbored variants from all three impact classes, indicating that a subset of genes accumulates multiple classes of mutation. These patterns were consistent across haplotypes and between SNVs and INDELs. For SNVs, missense variants were the most common moderate-impact class (∼4.3 per affected gene), whereas stop-gained variants predominated among high-impact SNVs. High-impact INDELs were more pervasive than high-impact SNVs, affecting 73% of INDEL-containing genes compared to 18% of SNV-containing genes, with frameshift variants as the dominant high-impact class (∼1.9 per affected gene) and disruptive in-frame insertions and deletions contributing primarily to the moderate-impact category (**Figure 5a-e**).

**Figure 5.**
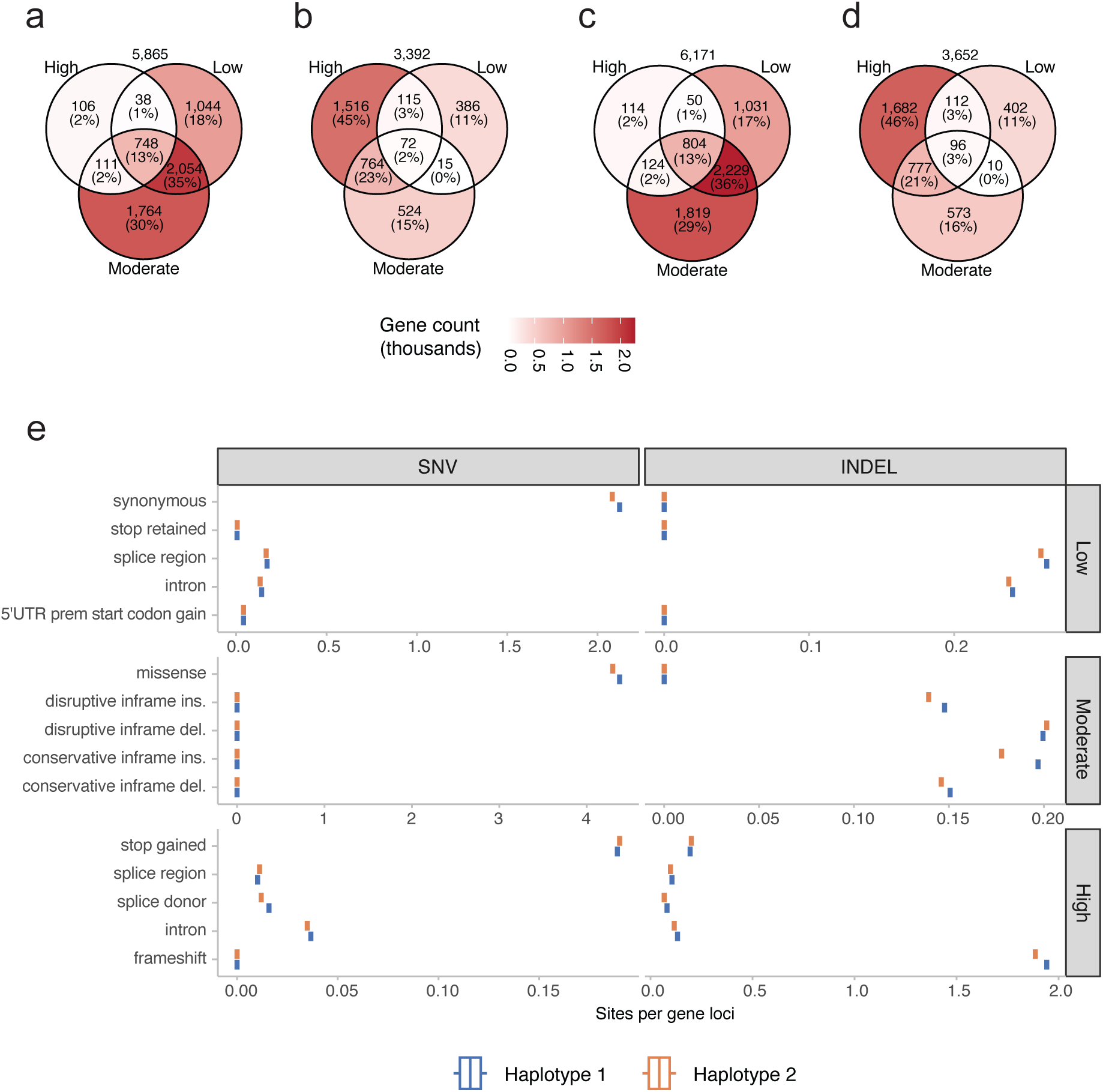
Distribution and functional impact of Carménère variants. Panels show the number and proportion of genes affected by variants in each predicted impact category for (a) SNVs in haplotype 1, (b) INDELs in haplotype 1, (c) SNVs in haplotype 2, and (d) INDELs in haplotype 2. Panel (e) shows the average number of variant sites per affected gene, grouped by variant type and predicted impact category. For each panel, proportions were calculated relative to the total number of genes affected by that variant type within the corresponding haplotype (total shown above each Venn diagram). Panels (a-d) share the same color legend.

### Functions of genes affected by clonal variants

Among genes affected by high-, moderate-, and low-impact variants, an average of 2,523 ± 85 genes per haplotype carried consensus heterozygous variants (i.e. variants present in all three replicates of at least one clone). Functional annotation using VitisNet (Grimplet et al., 2012) revealed representation across multiple biological pathways, with plant-specific signaling, secondary metabolism, transport, and signal transduction among the most affected categories (**Figure 6**; **Supplementary Tables 3 & 9**). Genes associated with plant-pathogen interaction networks were the most prominently affected across all impact categories and showed the largest gene counts relative to other pathways: on average per haplotype, 65, 163, and 121 genes belonging to the R-protein plant-pathogen interaction network carried high-, moderate-, and low-impact variants, respectively, corresponding to approximately 12%, 29%, and 22% of all annotated members of this network in the Carménère genome. Within secondary metabolism, genes in terpenoid, phenylpropanoid, and flavonoid biosynthesis were affected across all impact categories (**Figure 6**). Notably, genes involved in DNA replication and homologous recombination carried variants across all three impact levels, raising the possibility that somatic mutations in DNA repair pathways may themselves influence mutation accumulation rates in some clones.

**Figure 6.**
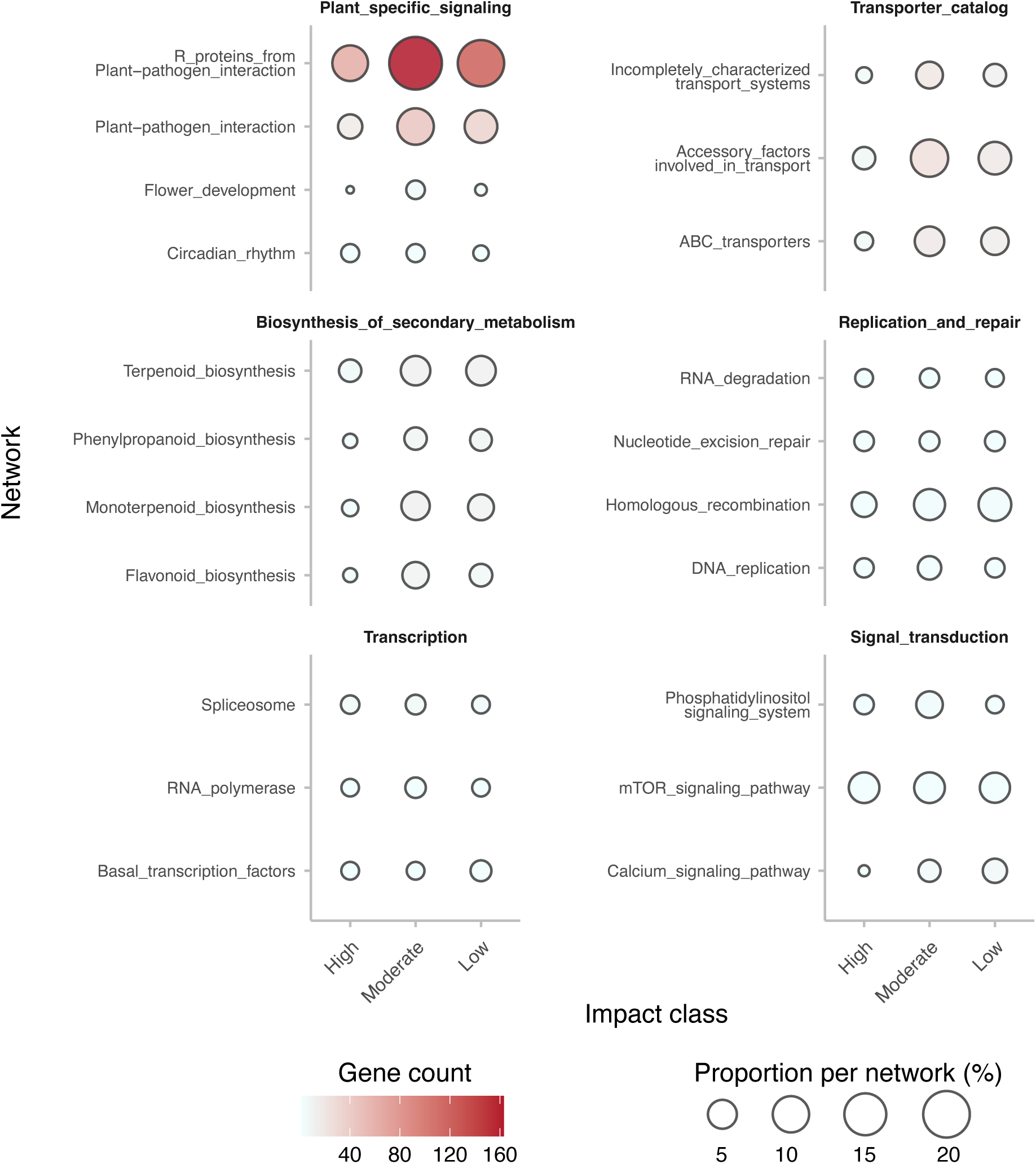
Functional networks of genes affected by consensus heterozygous variants across predicted impact classes. Bubble plot showing the number and proportion of genes per functional network affected by consensus heterozygous variants. For each impact class, the three general functional categories with the highest proportion of affected genes were selected. Within each general category, the top three networks with the highest numbers of affected genes are shown. Bubble size represents the number of affected genes per network, and color indicates the proportion relative to the total number of genes annotated in that network.

### Candidate genes associated with phenotypic variation among clones

To explore whether clonal variants might be associated to observed phenotypic differences, a variant phenotype consistency framework based on extreme phenotypes (Amanat et al., 2020) and gene-burden style carrier status (Floriani & Lipka, 2025) was applied to continuous traits measured across clones (as described in the methods section; **Supplementary Table 3**). For each trait, the three clones with the highest and lowest values were defined as extreme groups, and genes carrying high- or moderate-impact consensus heterozygous variants were screened for carrier patterns perfectly matching these groups. A total of 43 genes showed perfect consistency, 21 of which belonged to R-protein or plant-pathogen interaction networks (**Supplementary Table 10**). Among the remaining candidates, a gene encoding a pyruvate dehydrogenase E1 alpha subunit showed carrier status perfectly matching the three clones with the highest alcohol content after fermentation. A gene involved in auxin signaling was consistent with the three clones showing the highest anthocyanin levels and single berry weight. Two genes with phosphoglucomutase/phosphomannomutase functions showed carrier patterns consistent with clones exhibiting low anthocyanins, low pH, and high titratable acidity (**Supplementary Table 10**). Although the limited sample size precludes formal association testing, these results identify candidate genes and pathways potentially linked to phenotypic divergence among Carménère clones and provide a basis for targeted follow-up studies.

## Discussion

Traditional marker-based approaches for assessing clonal diversity in grapevine have limited resolution, often underestimating genomic variation and failing to distinguish clones that diverged through recent somatic mutations (Baneh et al., 2009; Imazio et al., 2002; Moncada & Hinrichsen, 2007; Zombardo et al., 2022). Whole-genome sequencing has substantially improved our ability to capture this variation. The first Carménère genome assembly represented an important step forward in the genomic characterization of this cultivar (Minio et al., 2019), but its fragmentation and partial phasing limited its utility for variant phasing, fine-scale analyses of genome structure, and clonal diversity studies. Here, we generated a high-quality phased diploid genome assembly of Carménère FPS 02, achieving over 99% BUSCO completeness, a substantial improvement over the previous version (∼93%). Most pseudomolecules correspond to full chromosomes, though a subset still lack telomeric repeats; accordingly, the assembly is best described as near-complete rather than telomere-to-telomere. The number and size of structural variants between the phased haplotypes are substantially reduced relative to the previous assembly, likely reflecting improved contiguity and more balanced haplotype representation rather than a true biological difference.

Kinship and PCA analyses confirmed clonal identity across samples while identifying Carolina77_RF20 as an outlier with consistently reduced relatedness to all other clones, suggesting a distinct genetic background or misclassification. This sample was excluded from downstream analyses. Across the remaining clones, most variants were shared among a large number of samples, indicating that a substantial fraction of observed variation reflects standing heterozygosity inherited from the ancestral genotype rather than recent clonal divergence. This pattern is consistent with Zinfandel, where 30% of heterozygous SNVs and 24% of heterozygous INDELs were shared across all 15 analyzed clones, while only 13% and 16% of heterozygous INDELs and SNVs, respectively, were private to individual clones (Vondras et al., 2019). To focus on variants more likely associated with recent clonal divergence (Callipo et al., 2025, 2026; Tello et al., 2025; Xiao et al., 2025), an allele frequency filter (AF ≤ 0.25) was applied, retaining approximately 3.4% of SNVs and 6.5% of INDELs per sample. Because this filter does not explicitly model clone structure, we complemented it with a replicate-aware consensus framework distinguishing fully supported clonal variants, partially fixed variants, and singletons.

Using this framework, we identified an average of more than 7,000 SNVs and 2,000 INDELs private to each clone, consistently supported across biological replicates. These values are consistent with the range of clone-private variants reported in other cultivars (Roach et al., 2018; Urra et al., 2023; Vondras et al., 2019), and reflect the expected accumulation of somatic mutations over repeated propagation cycles. While many of these mutations are likely neutral, evidence from cassava shows that deleterious mutations can become fixed during clonal propagation when recombination is limited and purging is ineffective (Ramu et al., 2017), and the functional impact of the variants identified here cannot be assumed negligible without further investigation. The number and consistency of these clone-private variants enable discrimination at higher resolution than traditional SSR or AFLP markers (Imazio et al., 2002; Moncada & Hinrichsen, 2007).

Direct numerical comparisons across studies are complicated by differences in reference genome quality, filtering strategies, and replicate structure, but the broader literature consistently supports the utility of genome-wide variation for resolving clonal structure. In Malbec, WGS of four clonal accessions identified 941 SNVs, of which 884 were clone-private; 41 validated markers were used to genotype 214 accessions, identifying divergent lineages associated with propagation history in Argentina versus France (Calderón et al., 2021). Malbec shares a direct historical parallel with Carménère: both were introduced to South America from France in the mid-to-late 19th century and propagated since in geographic isolation, suggesting that clonal lineages shaped by independent selection pressures could similarly be identified in Carménère if European accessions were included.

In Tempranillo, WGS of 35 plants from seven Iberian regions identified 1,120 SNVs across the samples with 158 considered highly-confident; a subset of 56 informative variants were used to genotype 185 plants, resolving three geographically structured lineages and tracing a westward dissemination route from the Ebro Valley (Tello et al., 2025). In Chardonnay, stringent k-mer-based filtering of variants from 15 clones yielded 1,620 markers that successfully discriminated clones and were validated on independently sourced material (Roach et al., 2018). A replicate-based approach, as used here and by Urra et al. (2023), reduces dependence on such stringency by supporting a larger and more reliable set of clone-private variants. Across 18 clones of Cabernet Sauvignon, Sauvignon Blanc, Chardonnay, and Merlot, approximately 4 million variants per clone were detected before filtering, comparable to the 4.2 million detected in Carménère, with up to 2,077 clone-private SNVs and 754 INDELs per clone identified after replicate-based filtering, a subset of which was validated by amplicon sequencing for clone discrimination (Urra et al., 2023). The larger number of clone-private variants in Carménère (∼9,000 SNVs and ∼2,000 INDELs) likely reflects differences in reference genome quality and filtering parameters rather than biology alone. The geographic structuring of clonal diversity observed in Malbec and Tempranillo raises the possibility that similar signals may be recoverable in Carménère; the Chilean clones studied here all derive from a single mid-19th century introduction from France, and including European accessions in future analyses would allow ancestral allele inference and potentially reconstruct the regional origins and dissemination history of the Chilean germplasm.

The higher density of variants in non-coding and repetitive regions is consistent with observations in Zinfandel and Malbec (Calderón et al., 2021; Vondras et al., 2019). Among variants affecting coding sequences, genes associated with plant-pathogen interactions were the most frequently impacted, a pattern well documented across plant genomes (Borevitz et al., 2007; Caicedo & Schaal, 2004; Karasov et al., 2014; Paineau et al., 2024; Wei et al., 2016). Resistance genes typically occur in clusters with high sequence similarity and repetitive content, features that promote diversification through replication slippage and non-allelic homologous recombination (Nagy & Bennetzen, 2008; Wicker et al., 2007). Variation was also observed in genes involved in secondary metabolism, pathways that influence both plant stress responses and the organoleptic properties of fruit and wine (Ali et al., 2010; Conde et al., 2007; Ferrandino et al., 2023; González et al., 2015; Murcia et al., 2017; Rienth et al., 2021).

Although the limited number of clones and sparse phenotypic data prevented formal association testing, an exploratory variant-phenotype consistency framework based on gene-burden logic and extreme-phenotype enrichment identified candidate genes whose variant patterns matched clones at the extremes of specific trait distributions (**Supplementary Table 10**). Given the number of genes tested relative to the number of clones, perfect consistency can occur by chance at non-negligible frequency; these results are strictly hypothesis-generating. Variants in a gene encoding a pyruvate dehydrogenase E1 alpha subunit were associated with higher alcohol content. This enzyme links glycolysis to the TCA cycle, and altered activity may influence carbon flux and fermentation-related substrate availability (Marillia et al., 2003; Yu et al., 2012), though this mechanistic interpretation remains speculative. Variants in genes with phosphoglucomutase/phosphomannomutase functions were associated with anthocyanin content, pH, and titratable acidity. These enzymes regulate carbohydrate partitioning, and reduced function has been associated with lower starch production and higher soluble sugar levels(Kofler et al., 2000; Malinova et al., 2014), which may affect organic acid and flavonoid synthesis (Malinova et al., 2014; Walker & Famiani, 2018; Zheng et al., 2009). Functional validation is required before any causal interpretation can be drawn.

Given the economic importance of Carménère in Chilean viticulture and its limited global distribution, characterizing its clonal diversity has direct implications for clonal selection and genetic resource conservation. Despite a narrow founding event and more than 150 years of uncontrolled vegetative propagation, substantial genomic diversity has accumulated among Chilean Carménère clones, largely invisible to earlier marker-based studies. While most detected variants likely represent neutral somatic mutations, a subset affects genes in metabolic and stress-response pathways that may influence agronomic and enological performance. Maintaining well-characterized germplasm collections that integrate genomic and phenotypic data will be essential for accurate clone identification, conservation, and selection. Future studies combining larger clonal panels, robust phenotyping, and functional validation will be needed to determine the biological significance of these variants and to better exploit clonal diversity for the improvement and long-term preservation of Carménère genetic resources.

## Data availability

The sequencing data generated for this project are available at NCBI (BioProject PRJNA1440530). The NCBI BioProject accession for sequencing data previously published and used in this study can be seen in **Supplementary Table 4**. Genome assemblies, gene models, repeats, variant files and Additional file 1 and 2 are publicly available at Zenodo (10.5281/zenodo.19139722). Dedicated genome browsers and BLAST tools for the Carménère genome are available at https://www.grapegenomics.com. A set of custom scripts used to process some data are deposited in GitHub at (https://github.com/jadgarci/Carmenere-diversity-supporting-scripts)

## Acknowledgements

The authors thank Viña Santa Carolina for providing access to the Bloque Herencia Carménère selections, for their long-term commitment to the conservation of Chilean pre-phylloxera grapevine genetic resources, and for their financial and logistical support of the field characterization program. The authors also thank the DNA Technologies Core Facility at the University of California, Davis, for sequencing support. This work was supported in part by the Ray Rossi Endowment in Viticulture and Enology.

## Supplementary Figures

**Supplementary Figure 1.**
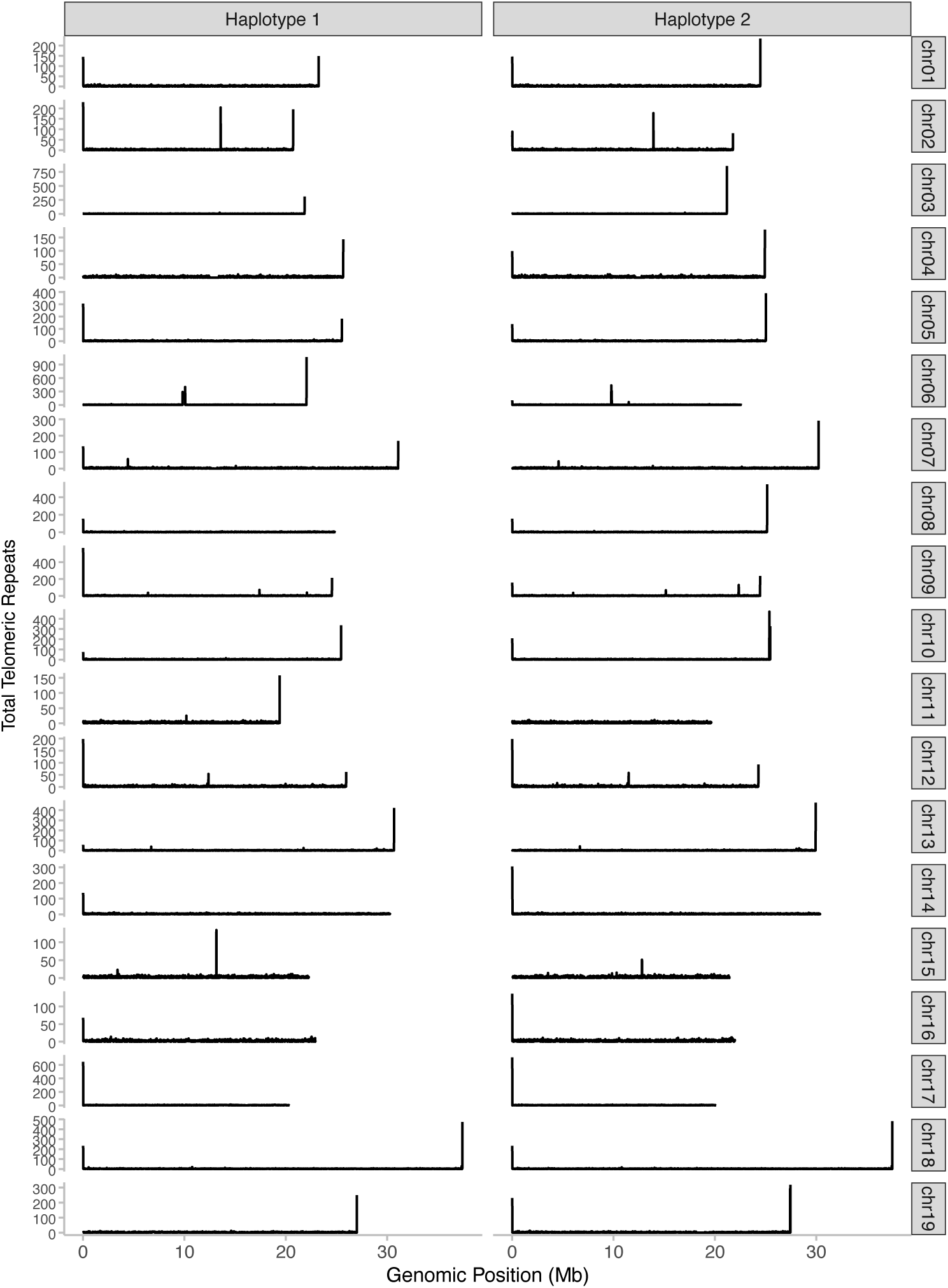
Distribution of telomeric repeat counts across pseudomolecules of the Carménère FPS 02 phased assembly. The grapevine telomeric repeat unit (TTTAGGG) was quantified in non-overlapping 10 kbp windows along each pseudomolecule. Counts are displayed separately for Haplotype 1 and Haplotype 2 to allow direct comparison of telomeric repeat distribution between haplotypes. Peaks at chromosomal extremities are indicative of telomeric regions.

**Supplementary Figure 2.**
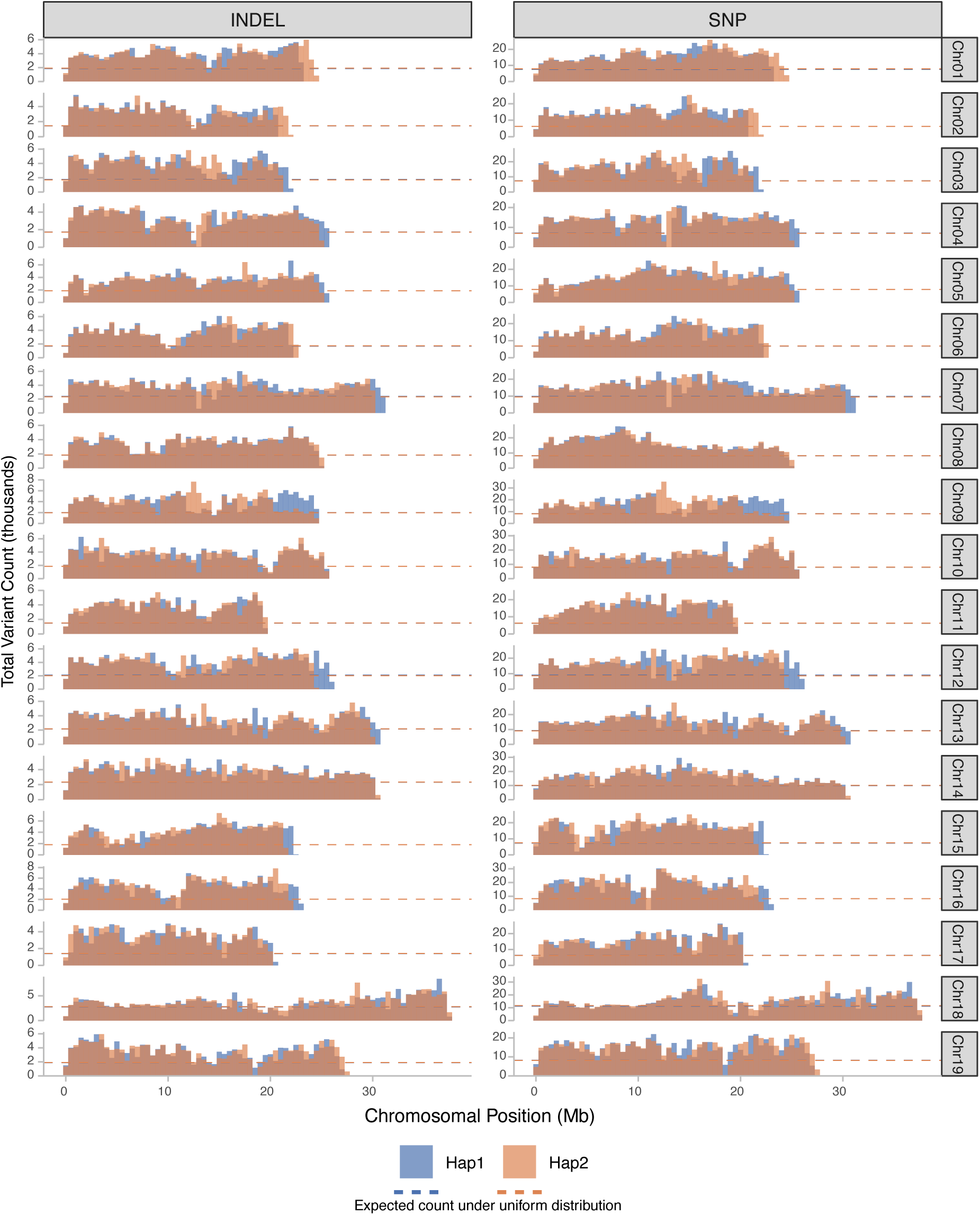
Variant density across pseudomolecules of the Carménère FPS 02 phased assembly. Single nucleotide variants (SNVs) and insertions/deletions (INDELs) were counted in non-overlapping 10 kbp windows and summarized as a histogram with 0.5 Mbp bins. Each panel corresponds to a pair of equivalent pseudomolecules from Haplotype 1 and Haplotype 2, overlaid for direct comparison. The dashed horizontal line indicates the expected variant count per bin under a uniform distribution across the pseudomolecule.

**Supplementary Figure 3.**
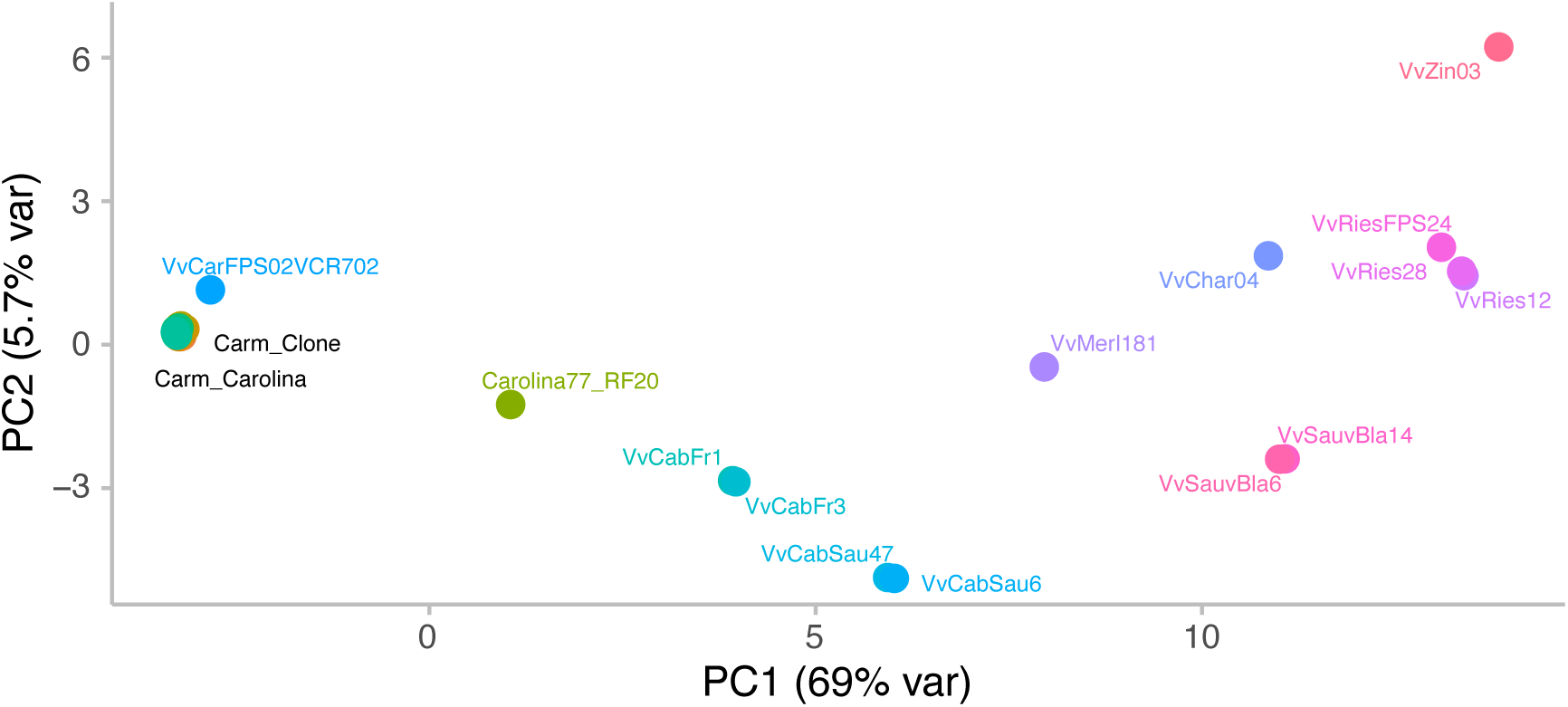
Principal component analysis of kinship-based genetic distances among Carménère clones and reference cultivars. Principal component analysis (PCA) of the pairwise distance matrix derived from kinship estimation across Carménère clones and outgroup cultivars. Each point represents one sample, coloured by sample. Abbreviations: Carm_Carolina, Carménère accessions named Carolina in the present study; Carm_Clone, Carménère accessions named Clone in the present study; VvCar, Carménère accession from a previously published dataset; VvCabFr, Cabernet Franc clones; VvCabSau, Cabernet Sauvignon clones; VvMerl, Merlot clone; VvSauvBla, Sauvignon Blanc clones; VvChar, Chardonnay clone; VvRies, Riesling clones; VvZin, Zinfandel clone.

**Supplementary Figure 4.**
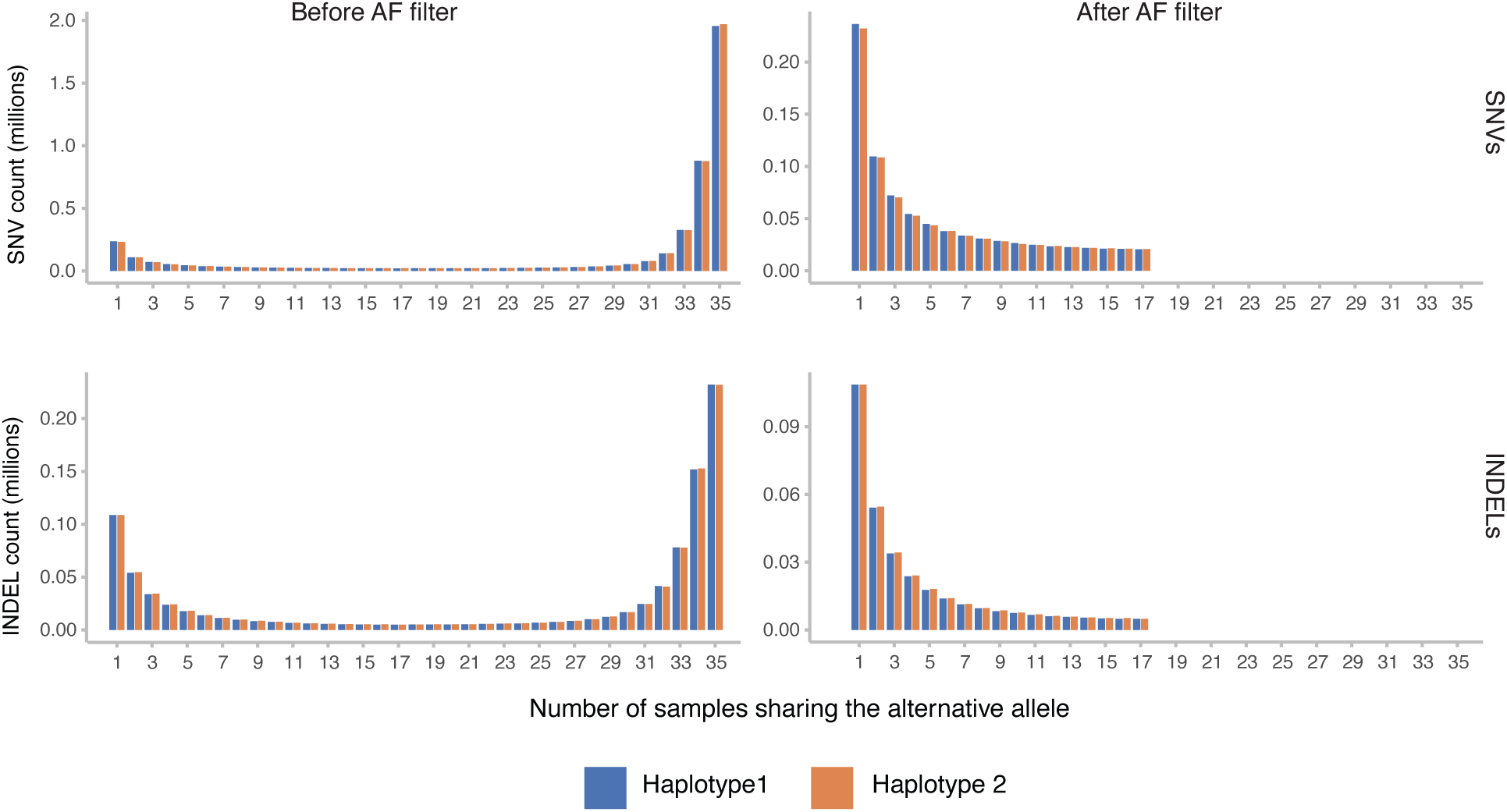
Distribution of variants by the number of samples sharing each variant, before and after alternative allele frequency filtering. Each bar represents the total number of variants (SNVs or INDELs) observed in exactly k samples, where k ranges from 1 to 35 samples. Distributions are shown before and after applying the alternative allele frequency filter (alternative allele frequency < 0.25).

**Supplementary Figure 5.**
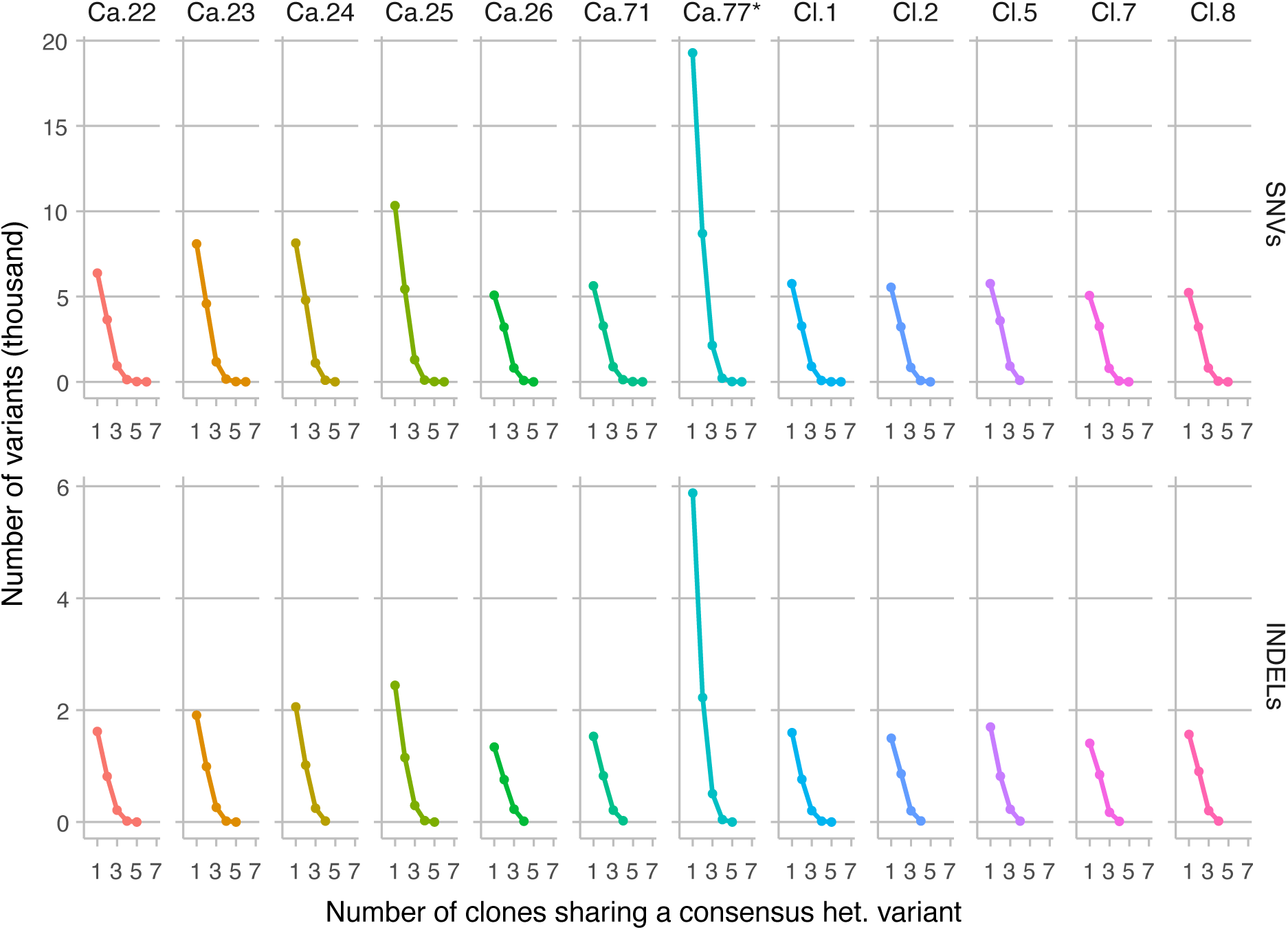
Per-clone contribution of SNVs and INDELs across sharing levels. Each panel displays the number of consensus heterozygous variants attributed to a given clone, broken down by sharing level k, where k denotes the number of clones carrying the variant. A consensus heterozygous variant was defined as a site heterozygous in at all replicates of a given clone. Name abbreviations are as follows: Ca., Carménère clones named Carolina in the study; Carménère clones named Clone in the study. *The elevated count of clone-private variants (k = 1) observed in Ca.77 reflects the fact that this clone was represented by only two replicates, such that consensus required agreement across two samples rather than three, lowering the stringency threshold relative to all other clones.

## Notes

### Competing Interest Statement

The authors have declared no competing interest.

